# Fear of the new? Geckos hesitate to attack novel prey, feed near objects and enter a novel space

**DOI:** 10.1101/2022.03.09.483578

**Authors:** Birgit Szabo, Eva Ringler

## Abstract

Neophobia, the fear of novelty, is an ecologically important response which enables animals to avoid potentially harmful situations. Low levels of neophobia have been linked to elevated dispersal/ migration, invasiveness and living in human modified landscapes albeit only in birds and mammals. In this study, we assessed neophobia in captive Tokay geckos (*Gekko gecko*). We expected to find low neophobia in our geckos because they are invasive and adopt well to anthropogenic environments. This species is, however, also both predator and prey in the wild which might select for higher neophobia. We tested neophobia in three contexts: attacking novel prey, foraging near novel objects and entering a novel space. We aimed to quantify (1) neophobia in these contexts, (2) individual consistency across trials using different novel stimuli, and (3) correlation of individual responses across the three contexts. We found that geckos hesitate to attack novel prey and prey close to objects (familiar and novel). Geckos hesitated the most when entering novel space and repeatability of behaviour across contexts was low (R = 0.12) indicating that neophobia might not be a single trait. The strength of a neophobic response can indicate how anxious or curious an individual is. This test has great potential to help answer questions about how captivity, enrichment, rearing environment and cognition affect fear responses in different contexts in lizards. By studying reptiles, we can better understand the universality of what is known about the causes leading to difference in neophobia across individuals and species.

## Introduction

Neophobia is the fear of novelty and is expressed by hesitation to approach or complete avoidance of a novel stimulus that has never been encountered before (Crane, Brown, Chivers, & Ferrari, 2020; Greenberg & Mettke-Hofmann, 2001). The recognition of novelty, as opposed to familiarity, is a cognitive process and is transitive in its nature based on the experience of an individual (Greenberg & Mettke-Hofmann, 2001). In the brain, a novel stimulus activates a large neural network that enhances attention but after repeated exposure neural activity in response to the stimulus decreases (Ranganath & Rainer, 2003). Novel stimuli can elicit varying degrees of neophobic reactions if, for example, a novel stimulus is innately recognised because avoidance behaviour is conserved across populations, species or higher taxa (Burghardt, 1973; Saul & Jeschke, 2015). Furthermore, a novel stimulus might be recognised due to its similarity with known stimuli through generalisation (Crane et al., 2020). The background risk level also plays a major role in how intensely an individual reacts to a novel stimulus (Greenberg & Mettke-Hofmann, 2001; Ranganath & Rainer, 2003; Vernelli, 2014). Low levels of risk or uncertainty (evolutionary: the environment a species evolved in; individual: the past experience, i.e. learning) can decrease neophobia (e.g. Barnett, 1958; Mitchell, 1976) while high levels can increase neophobia (e.g. Brown, Ferrari, Elvidge, Ramnarine, & Chivers 2013; Brown, Elvidge, Ramnarine, Ferrari, & Chivers, 2015). The response to a novel stimulus depends, therefore, on the evolutionary background of a species, past experience and cognitive skill of an individual and the context in which the novel stimulus is encountered (Crane et al., 2020; Ranganath & Rainer, 2003).

Neophobia is an ecologically important response to avoid potentially harmful situations or individuals in their environment (Crane et al., 2020; Greenberg & Mettke-Hofmann, 2001), and determines an individuals’ likelihood to approach a novel stimulus or to avoid it (Crane et al., 2020; Greenberg & Mettke-Hofmann, 2001; Lima & Dill, 1990). If a threat is real and avoided, an individual can escape harm (= correct response; Lima & Dill, 1990). If, however, an individual does not avoid a true threat then it might pay a large cost such as physical harm or even death (= false negative). The opposite, neophobia directed at a harmless stimulus (= false positive), has a low cost (missed opportunity) and explains why animals generally err on the side of caution (Crane & Ferrari, 2017; Crane et al., 2020). Consequently, to minimise costs, neophobic responses should be relatively plastic (Brown et al., 2013; Crane & Ferrari, 2017; Crane et al., 2020).

Neophobia is most often explored in three contexts: When encountering new food, new objects and new space (Crane et al., 2020; Greggor, Thornton, & Clayton, 2015). Food neophobia is measured by presenting an individual with a novel food within a familiar environment. The time taken to consume the novel food is recorded. To properly assess the presence of neophobia, responses are compared to those towards familiar food (Greggor et al., 2015). To test object neophobia, individuals are confronted with a novel object close to familiar food within a familiar environment and the latency to eat compared to the control without the novel object is recorded (Greggor et al., 2015; Takola, Krause, Müller, & Schielzeth, 2021). Testing food and object neophobia within a familiar environment is important, because background familiarity (context) can affect the intensity of a neophobic reaction. Typically, higher levels of neophobia are expressed in a familiar rather than an unfamiliar environment (Greenberg & Mettke-Hofmann, 2001; Ranganath & Rainer, 2003; Vernelli, 2014). Finally, space neophobia is assessed by measuring the time taken to enter a novel environment (Greggor et al., 2015). Novelty is, of course, not just restricted to the visual domain. Novel smells or sounds can be important sources of information but these are rarely considered except in studies directly looking at predator neophobia (Crane & Ferrari, 2017; Crane et al., 2020).

Differences in neophobia are found within and across species. Research (mostly object and food neophobia) revealed that juveniles show lower neophobia compared to adults (e.g. birds: *Milvago chimango*, Biondi, Bo, & Vassallo, 2010; Guido, Biondi, Vasallo, & Muzio, 2017; mammals: *Cebus apella*, Visalberghi, Janson, & Agostini, 2003), but it rarely differs across the sexes (e.g. birds: *Passer domesticus*, Ensminger, Westneat, & Zeh, 2012; mammals: 10 ungulate species, Schaffer et al., 2021; but see Crane & Ferrari, 2017 and Crane et al., 2020 for a short discussion). Furthermore, neophobia can be influenced by the social environment (e.g. birds: 10 corvid species, Miller et al., 2022; *Taeniopygia guttata*, St. Lawrence, Rojas Ferrer, & Morand-Ferron, 2021; mammals: *Bos taurus taurus*; Meagher et al., 2015; 10 ungulate species, Schaffer et al., 2021) and might negatively influence other cognitive processes such as learning (e.g. birds: *Milvago chimango,* Guido et al., 2017) but not all studies found such a link (e.g. fish: *Neolamprologus pulcher,* Bannier, Tebbich, & Taborsky, 2017; lizards: *Podarcis erhardii,* De Meester, Pafilis, & Van Damme, 2022). Finally, species differences in neophobia can be explained by the trophic level they inhabit (Crane & Ferrari, 2017), diet (Mettke-Hofmann, Winkler, & Leisler, 2002), habitat urbanisation (Miller et al., 2022) and their tendency to exploit new habitats (invasiveness) or migrate (Greenberg & Mettke-Hofmann, 2001). Overall, neophobia is studied across vertebrates but research in lizards is scarce (Crane et al., 2020). A study in the Aegean wall lizard (*Podarcis erhardii*) tested object neophobia by measuring the time it took lizards to consume food next to an unfamiliar object (De Meester et al., 2022). Other studies in lizards focused on neophilia, the attraction towards novelty (Takola et al., 2021). For example, object neophilia was tested in the Italian wall lizard (*P. siculus*) and the Geniez’s wall lizard *(P. virescens)* to quantify boldness. Object neophilia was also measured towards a novel cylinder to investigate its’ effect on inhibitory control in five skink species (Szabo, Noble, & Whiting, 2019; Szabo, Hoefer, & Whiting, 2020). Attraction to novel food as part of a personality assay was tested in the water skink (*Eulamprus quoyii*; Carazo, Noble, Chandrasoma, & Whiting, 2014) and exploration of novel space and object neophilia were quantified in the leopard gecko (*Eublepharis macularius*) to investigate novelty recognition (Kundey & Phillips, 2021). Lastly, bearded dragons (Pogona vitticeps) latency to start moving in a novel environment was measured to investigate if such tests can be used to evaluate welfare (Moszuti, Wilkinson, & Burman, 2017). Lizards are extremely diverse regarding their ecology, life-history, and behaviour (Pianka, & Vitt, 2003) which makes them excellent models to investigate the relationship of neophobia to a range of traits. By investigating neophobia in lizards we will be able to better understand the universality of what is known about the causes leading to difference in neophobia across individuals and species.

In this study, we assessed neophobia in three contexts: attacking novel prey, foraging near novel objects and entering a novel environment in captive tokay geckos (*Gekko gecko*). Tokay geckos are a medium sized, nocturnal, arboreal, insectivorous lizard species from South-East Asia which form temporary family groups (Grossmann, 2006). Our aim was to (1) quantify neophobia and potential effects of body condition, sex and temperature, to (2) quantify consistency across trials using different novel stimuli, and (3) to assess if responses are correlated across contexts (repeatability). Tokay geckos are an excellent species to test neophobia across different contexts. In their natural habitat they are both predator and prey (Grossmann, 2006) and might therefore show both food and object neophobia (Greenberg & Mettke-Hofmann, 2001; Mettke-Hofmann, Winkler, Hamel, & Greenberg, 2013). Furthermore, they express a range of anti-predator behaviours including freezing, flight, defensive displays (mouth gaping) and defensive barks associated with feigned attacks in captivity as well as in the wild (Grossmann, 2006). Such a large range of anti-predator behaviour could be a result of high background risk in their natural environment resulting in the evolution of neophobic responses to a range of stimuli. These lizards are also invasive and very successful inhabitants of urbanised landscapes (Grossmann, 2006; Rocha, Piva, Batista, & Coutinho Machado, 2015) which has been linked to low levels of neophobia (Greenberg & Mettke-Hofmann, 2001; Miller et al., 2022). Robustly quantifying neophobia in our captive geckos will open up new avenues for future research into how anxiety is related to welfare, cognitive ability, personality/coping style and invasiveness in lizards.

## Methods

### Animals, captive conditions and husbandry

22 captive bred, adult, naïve tokay geckos, 10 males (SVL range = 11.35-15.02 cm) and 12 females (SVL range = 11.29-13.72 cm) (Grossmann, 2006) approximately 2-6 years of age from different breeders were used in this study. Sex was determined by the presence (male) and absence (female) of femoral glands (Grossmann, 2006). Geckos are kept singly in plastic terraria (females – 45 L x 45 B x 70 H cm; males – 90 L x 45 B x 100 H cm) equipped with a compressed cork back wall, cork branches, refuges made out of cork branches cut in half hung on the back wall and life plants. Additionally, enclosures are equipped with a light on top to provide lizards with UVB (Exo Terra Reptile UVB 100, 25 W). Enclosures contain a drainage layer of expanded clay with organic rainforest soil (Dragon BIO-Ground) on top. The layers are separated by a mosquito mesh to prevent mixing. On top of the soil we spread autoclaved red oak leaves. Our enclosures are bio-active with collembola, isopods and earth worms in the soil that break down the faecal matter produced by the geckos.

Enclosures are set up on shelfs across two rooms. Small enclosures are kept on top of the shelves and large enclosures on the bottom. We keep lizards in a fully controlled environment with a reversed 12h:12h photo period (light: 6pm to 6am, dark: 6am to 6pm) to accommodate their nocturnal lifestyle. A red light (PHILIPS TL-D 36W/15 RED) not visible to geckos (Loew, 1994) is kept on 24h a day and enables researchers to work with the animals during their active period. Sunrise and sunset are simulated automatically accompanied by a gradual change in temperature which reaches approximately 25 °C during the night cycle and 30 °C during the day cycle. A heat mat (TropicShop) is fixed to the outside of each enclosure increasing the temperature by 4-5 °C for thermoregulation. To mimic natural tropical conditions, the humidity is kept at 50% and daily rainfall (osmotic water, 30s every 12h at 5pm and 4am) increases the humidity to 100% for a short period of time.

Lizards are fed 3-5 adult house crickets (*Acheta domesticus*) three times per week on Monday, Wednesday and Friday. Crickets are gut loaded with cricket mix (reptile planet LDT), Purina Beyond Nature’s Protein™ Adult dry cat food and fresh carrots to ensure that they provided optimal nutrition (Vitamin D and calcium). We feed lizards with 20 cm long forceps to monitor their food intake. Water is provided *ad libitum* in a water bowl. To keep track of our lizards’ health, we weigh them once a month and measure their snout vent length every two months.

### Food and object neophobia

#### Testing procedure

Lizards were tested in their home enclosures to reduce stress of handling (Langkilde & Shine, 2006) and ensure strong neophobic responses (Greenberg & Mettke-Hofmann, 2001; Vernelli, 2014). At the start of a session, we placed a dim white light (LED, SPYLUX^®^ LEDVANCE 3000K, 0.3 W, 17 lm) on top of the tank. Lizards were trained to expect food or testing when this light was placed on top of their tank. Next, a lizard was located and if under a refuge the refuge was gently removed to expose the lizard. Then, the experimenter presented the stimuli in 20 cm long forceps within 4-5 cm of the lizard’s snout for a maximum of one minute. This distance was chosen as it represented the optimal attack distance (personal observation). Lizards were tested with familiar and novel foods, coloured and natural, to test food neophobia and dietary conservatism and with familiar and unfamiliar objects in a feeding context to test object neophobia (see detailed description below). One session was given per day, the inter-session interval was four days and each test was repeated twice with 14 days in between using new novel food and a novel object. The order in which lizards were tested in each test was randomised but counterbalanced. Lizards first tested on food neophobia and then object neophobia in the first repetition were tested on object neophobia and then food neophobia in the second repetition. The order in which lizards were tested within a given session was randomised to account for order effects. Trials were recoded using a Samsung S20 smartphone (108 Megapixel, 8K-FUHD). Testing was done between the 6^th^ and 25^th^ of October 2021 between 8:00 and 10:15 am.

#### Food neophobia and dietary conservatism

We used adult house crickets (*Acheta domesticus*) as familiar food. In the first repetition, we presented *Tenebrio molitor* larvae approximately last instar (Park et al., 2014) as the unfamiliar food and in the second repetition we presented desert locusts (*Schistocerca gregaria*) approximately 4^th^ immature instar (Samejo & Sultana, 2019) as the unfamiliar food. Geckos had no experience with these prey items for at least one year. Lizards were presented with four trials per session in a random order: (1) familiar cricket uncoloured, (2) familiar cricket coloured, (3) unfamiliar food uncoloured, (4) unfamiliar food coloured (electronic supplementary material, video M1). A trial lasted for a maximum of 60 seconds. To tease apart food neophobia from dietary conservatism we presented coloured and uncoloured, familiar and unfamiliar foods. We coloured insects using black (Fun Cakes, Colour dust, black, E153) or white (Fun Cakes, Colour dust, white snow, E171) powder food colouring (Figure 1, A-C).

**Figure 1.**
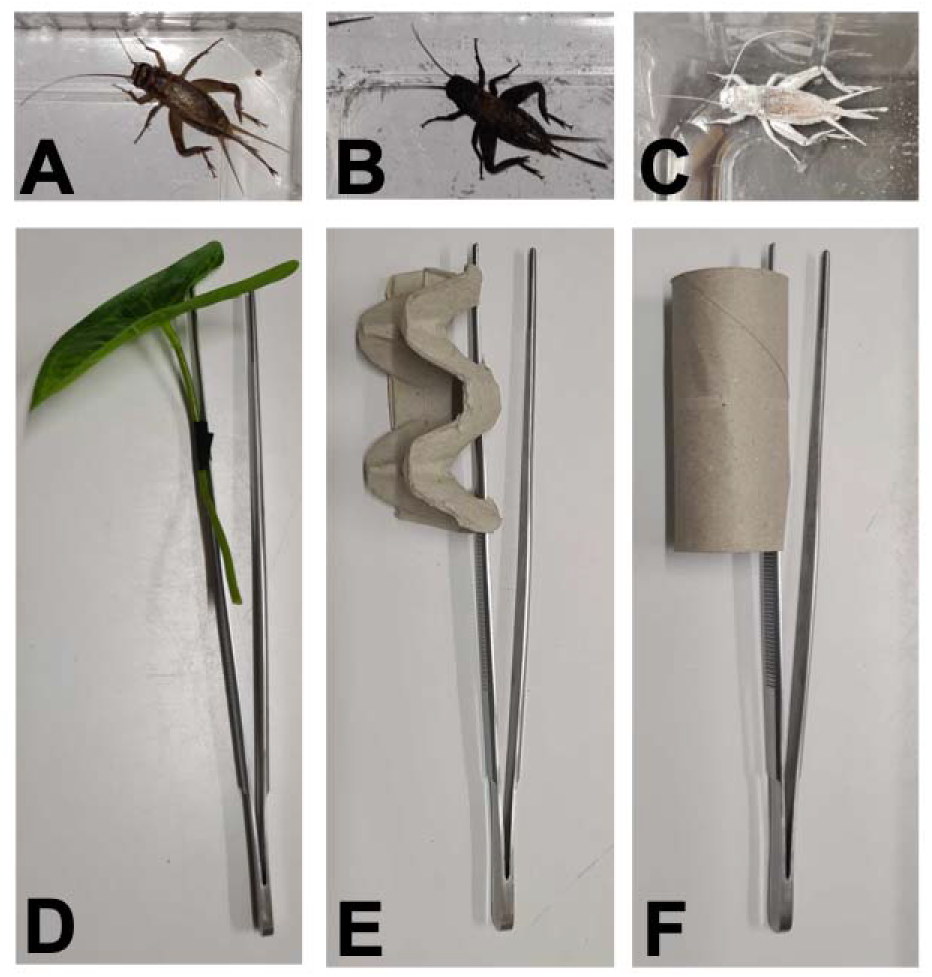
Examples of stimuli used in the food and object neophobia tests. A-C) An adult female cricket uncoloured (= natural, A), coloured black (B) and coloured white (C). D-E) The familiar artificial leaf (D) and the novel objects made out of cardboard (egg carton, E, and toilet paper roll, F).

#### Object neophobia

As the familiar object we used a plastic leaf that had been introduced for five days before testing into each lizard’s enclosure. As novel objects we used a piece of egg carton cut to size (9.5 cm L, 4.5 cm H, 4 cm W) in the first repetition and a cardboard toilet paper roll (9.5 cm L, 4 cm diameter) in the second repetition. Both novel objects were of similar size, material and colour but different in shape and were unfamiliar to lizards. Familiar and unfamiliar objects were attached to 20 cm long forceps (Figure 1, D-F) in the exact same position. Each object was only used once and each individual was tested with a new object. Lizards were presented with three trials per session in a random order: (1) familiar cricket (control), (2) familiar cricket next to the familiar object, (3) familiar cricket next to the unfamiliar object (electronic supplementary material, video M1).

### Space neophobia and object neophilia

#### Testing setup

Lizards were tested in empty glass testing tanks (45 L x 45 B x 60 H cm, ExoTerra) with three sides covered on the outside with black plastic to make them opaque (Figure 2, A). One testing tank was placed on a table in each animal room at 100 cm distance facing (with the front transparent doors) a wall. A dim white light (LED, SPYLUX^®^ LEDVANCE 3000K, 0.3 W, 17 lm) was placed in the top right corner of the terrarium and a camera (GoPro, Hero 8; linear mode, 1080 resolution, 24 FPS) mounted on a tripod recorded trials from above at 40 cm distance from the terrarium mesh lid (Figure 2, A). Trials lasted for 20 minutes. We first tested space neophobia directly followed by object neophilia resulting in 40 minutes for the whole test.

**Figure 2.**
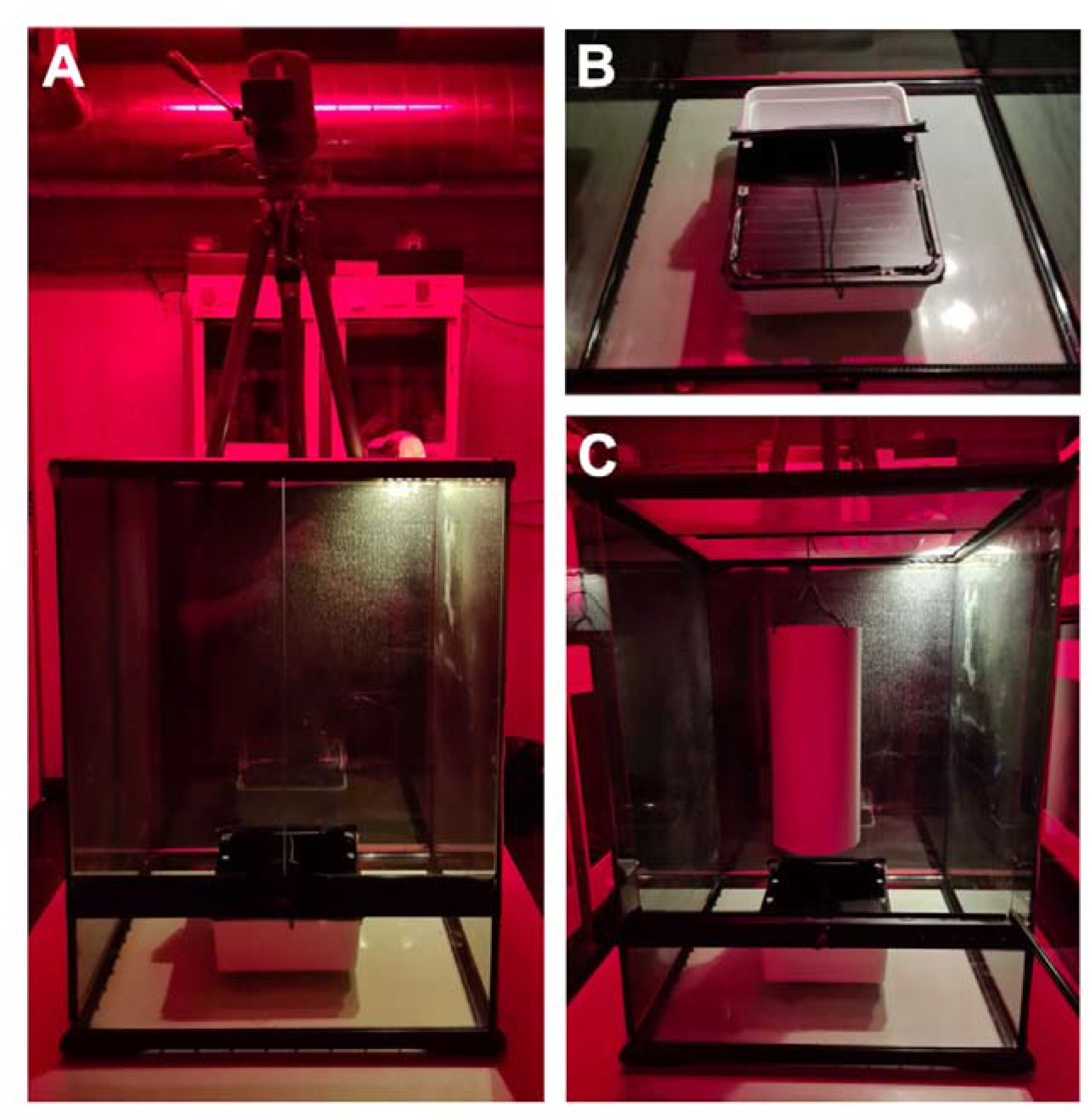
Picture of the glass testing tank, opaque box and PCV tube used during the space neophobia and object neophilia test. A) Experimental setup with the glass tank (45 L x 45 B x 60 H cm) wrapped on three sides in black plastic (doors closed), the dim white light (LED, SPYLUX^®^ LEDVANCE 3000K, 0.3 W, 17 lm) in the right, back corner of the lid, the camera (GoPro, Hero 8) fixed to a tripod filming from above and the opaque box (lid open) placed in the center of the testing tank. (B) Opaque box with the lid fixed to stay open (wire) in the middle of the glass tank. (C) Testing tank with the PVC tube hung above the open opaque box as done during the object neophilia test.

#### Testing procedure

First, a lizard was captured in an opaque, plastic box (24 cm L x 18 cm W x 7.5 cm H; white opaque bottom with a lid covered in black isolation tape; lids included 6 air holes). After capture, the individual was slowly carried to the testing terrarium and the box placed inside (ground, center, with the opening facing the back wall). Lizards were left alone for 5 minutes to calm down. Next, the experimenter started the video recording and opened a third of the box lid carefully. Using a wire, the lid was secured to stay open (space neophobia test) and provide an exit for lizards (Figure 2, B). Thereafter, the door to the testing terrarium was closed and the experimenter left the room. Lizards were left undisturbed for 20 minutes (electronic supplementary material, video M1). Then, the experimenter returned and carefully placed a light grey PVC tube (30 cm L, 12.4 cm diameter, 1 mm thick PVC) approximately 5 cm above the plastic box inside the testing terrarium by hanging it from the mesh ceiling with a wire (object neophilia test, Figure 2, C). Again, the test terrarium doors were closed and lizards were left undisturbed for another 20 minutes (electronic supplementary material, video M1). At the end of the object neophilia test, the lizard was recaptured either in the opaque box (when still inside) or a transparent box (allowing easier capture) and released back into its home enclosure. Afterwards, the testing terraria, opaque box and the PVC tube were thoroughly cleaned with 70% ethanol to remove the scent of the previous test subject. Equipment was left for 15 minutes for the alcohol to vanish. Each lizard was only tested once for space neophobia and object neophilia. Tests were conducted between 8:15 am and 14:45 am (active period of our lizards). All individuals were tested across two days (Tuesday and Thursday, none feeding days) within the same week on the 14th and 16^th^ of December 2021.

### Data collection

#### Food and object neophobia

We measured the time from when the lizard first noticed a food item until the first strike regardless of if the food was captured or not (capture latency) and from when an individual focused on a prey item the last time before striking (strike latency). We assumed that a food item was first noticed when a lizard moved its’ head to focus on it (on rare occasions only the eyes moved not the whole head). We used the free behavioural coding software BORIS (Friard & Gamba, 2016) to measure latencies to an accuracy of 0.001 seconds. To this end, videos were slowed down to half their speed. If no attack occurred, we recoded occurrence as 0 and assigned this data point a latency of 60 seconds. Videos were scored twice and latency measures across the two scores were highly consistent (Spearman rank correlation, r_s_ = 0.883, S = 555298, *p*-value < 0.001).

#### Space neophobia and object neophilia

From the trial videos we scored at what time in the trial (exit latency, in seconds) a lizard exited the opaque box by lifting its’ tail base over the rim of the box (= exiting with their whole body not counting the tail). If a lizard did not exit the box, we assigned it a latency of 1200 seconds (= 20 minutes). We also scored if the lizard touched the cylinder hung in the middle of the empty testing enclosure (yes – 1, no – 0, Bernoulli variable) and at what time in the trial (touch latency, in seconds).

### Statistical analyses

First, we wanted to know if lizards time to attack prey was influenced by its novelty (new prey species or novel colour) or an object (familiar or novel) presented close to the prey. Furthermore, we were interested if capture and strike latency were appropriate measures of neophobia. To this end, we analysed each test (food, object and space) separately running two models one with the log-transformed capture latency as the response variable and one with log-transformed strike latency (food and object neophobia only). Latencies were log transformed to conform to assumptions of normality. We used Bayesian generalised linear mixed models (MCMCglmm package, Hadfield, 2010) with a common weak prior for gaussian data because we had no prior knowledge of how our lizards would perform in this test. To analyse food neophobia and dietary conservatism we used stimulus (novel or familiar) and colour (natural or coloured) as two of the fixed effects. To analyse object neophobia we used stimulus (control, novel or familiar) as one of the fixed effects. Additionally, we were interested if trial (repetition one and two), sex (male or female), room (room 2 or room 5) or tank size (small or large) affected either measure. Moreover, whenever possible, we also added presentation order, test order, temperature and body condition (scaled mass index, Peig & Green, 2009, for details see the electronic supplementary material) as non-categorical fixed effects.

We were also interested which test elicited the stronger response by only looking at responses to novelty (food neophobia: response to novel food regardless of colour, object neophobia: response to both objects separately and space neophobia). To make latency measures comparable across tests we calculated relative latency. We divided latency measured in the food and object neophobia (capture latency) test by 60 seconds (= maximum trial length) and the latency to exit into a novel environment by 1200 seconds (= maximum trial length). We ran a MCMCglmm with the log-transformed relative latency as the response variable and stage (food, object and space) as the fixed effect.

Finally, we wanted to know if our lizards’ responses were correlated indicating a consistent trait and if responses were repeatable within individuals and across contexts. To this end, we performed a Spearman rank correlation test on the responses to novelty only to investigate if average measures of latency (capture and exit latency) correlated across tests (adjusted alpha = 0.016 [alpha 0.05/3] due to repeated analysis). We also calculated adjusted repeatability accounting for test and stimulus using the package rptR (Stoffel, Nakagawa, Schielzeth, & Goslee, 2017) for capture and exit latency. Finally, we plotted each individual’s response to novelty across contexts to visually look for individual differences.

All MCMCglmm included a random intercept of animal identity and a random slope of trial nested in session to account for non-independence and autocorrelation across successive choices due to repeated measures of trial and session across individuals. In the case of the space neophobia, the random effect only included animal identity because we only tested one trial. In all cases, we ensured that lags did not correlate (< 0.1; no auto-correlation; Hadfield, 2010), that the MCMC chain mixed sufficiently (by visually inspecting plots; Hadfield, 2010) and we performed a Heidelberg and Welch diagnostic tests to confirm that the MCMC chain was run for long enough (Hadfield, 2010). All analyses were conducted in R version 4.0.3 (R Core Team, 2020). We report our results based on the guidelines proposed by Muff and colleagues (2022): *p* > 0.1 no evidence, 0.1 < *p* < 0.05 weak evidence, 0.05 < *p* < 0.01 moderate evidence, 0.01 < *p* < 0.001 strong evidence, *p* < 0.001 very strong evidence. We only report results with at least moderate evidence within the text but provide all results in the electronic supplementary material. All raw data sets and code for analysis are available on the Open Science Framework (link for review purposes: https://osf.io/fhw64/?view_only=bf8bef37ef7e4f55beec48706adaa82a).

### Ethical note

We followed the guidelines provided by the Association for the Study of Animal Behaviour/ Animal Behaviour Society for the treatment of animals in behavioural research and Teaching (2022). All testing was approved by the Suisse Federal Food Safety and Veterinary Office (National No. 33232, Cantonal No. BE144/2020). Captive conditions were approved by the Suisse Federal Food Safety and Veterinary Office (Laboratory animal husbandry license: No. BE4/11).

## Results

### Food neophobia

We found very strong evidence that lizards hesitated to attack novel prey (MCMCglmm, estimate = 0.625, CI_low_ = 0.293, CI_up_ = 0.951, *p*-value = 0.000134) regardless of colour (MCMCglmm, estimate = 0.044, CI_low_ = −0.286, CI_up_ = 0.370, *p*-value = 0.790117) when we analysed capture latency (Figure 3). When looking at strike latency, we found similar strong evidence that lizards hesitated to attack novel prey (MCMCglmm, estimate = 0.503, CIlow = 0.210, CI_up_ = 0.796, *p*-value = 0.00187) regardless of colour (MCMCglmm, estimate = 0.255, CI_low_ = −0.038, CI_up_ = 0.545, *p*-value = 0.08801; electronic supplementary material, Figure S1). We found moderate evidence that capture latency was shorter in lizards from room 5 (MCMCglmm, estimate = −0.593, CI_low_ = −1.061, CI_up_ = −0.095, *p*-value = 0.021903; electronic supplementary material, Figure S2), and we found moderate evidence for longer strike latency later in the day (MCMCglmm, estimate = 0.049, CI_low_ = 0.010, CI_up_ = 0.088, *p*-value = 0.01549, electronic supplementary material, Figure S3). Finally, trial had no effect on response time indicating that both novel foods were of similar novelty (MCMCglmm, estimate = 0.021, CI_low_ = −0.443, CI_up_ = 0.544, *p*-value = 0.936361, electronic supplementary material, Table S1). None of the other fixed effects were affecting latency (electronic supplementary material, Table S1 and S2).

**Figure 3.**
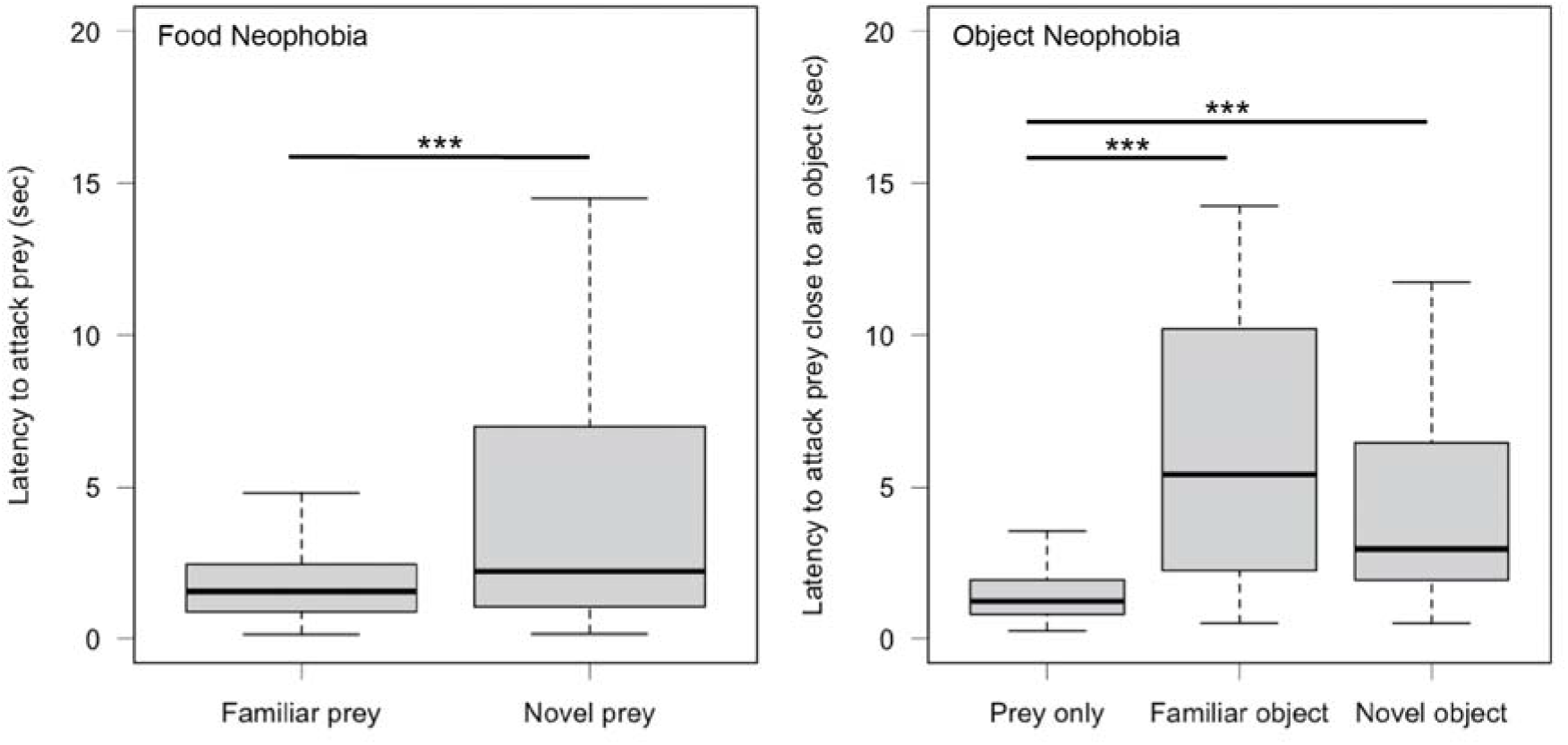
Box plots showing the capture latency to attack familiar (cricket) and novel food (mealworm or locust) in the food neophobia test and prey only, prey next to a familiar object (artificial leaf) and a novel object (egg carton or toilet paper roll) in the object neophobia test. The bold line indicates the median, the upper edge of the box represents the upper quartile, the lower edge the lower quartile, the top edge of the whisker the maximum and the bottom edge of the whisker the minimum (outliers are not shown). *** *p* < 0.001.

### Object neophobia

Our analyses revealed very strong evidence that lizards attacked prey close to objects more slowly than control prey regardless of familiarity both when capture latency (MCMCglmm, novel: estimate = 1.138, CI_low_ = 0.732, CI_up_ = 1.527, *p*-value < 0.00004; familiar: estimate = 1.456, CI_low_ = 1.050, CI_up_ = 1.847, *p*-value < 0.00004, Figure 3) and strike latency were analysed (MCMCglmm, novel: estimate = 0.683, CI_low_ = 0.313, CI_up_ = 1.043, *p*-value = 0.000464; familiar: estimate = 0.656, CI_low_ = 0.282, CI_up_ = 1.024, *p*-value = 0.000437, electronic supplementary material, Figure S1). We also found strong evidence for longer strike latency from lizards in small tanks (MCMCglmm, estimate = 0.971, CI_low_ = 0.303, CI_up_ = 1.639, *p*-value = 0.005096; electronic supplementary material, Figure S3). Finally, trial had no effect on latency indicating that both novel objects were of similar novelty (MCMCglmm, estimate = −0.189, CI_low_ = −0.645, CI_up_ = 0.260, *p*-value = 0.4081, electronic supplementary material, Table S3). None of the other fixed effects were influencing our measurements (electronic supplementary material, Table S3 and S4).

### Space neophobia and object neophilia

Our analysis revealed no effect of any of the tested variables on latency to exit a box into a novel environment (electronic supplementary material, Table S5). Furthermore, none of the lizards touched the novel object during the 20 minutes of the object neophilia test.

### Cross context analyses and repeatability

We found very strong evidence that lizards showed less neophobia when attacking novel prey (relative latency: MCMCglmm, estimate = −2.441, CI_low_ = −3.012, CI_up_ = −1.833, *p*-value < 0.00007) and when attacking prey near novel objects (MCMCglmm, estimate = −2.158, CI_low_ = −2.800, CI_up_ = −1.517, *p*-value < 0.00007) compared to when exiting into a novel environment (MCMCglmm, estimate_intercept_ = −0.581, CI_low_ = −1.147, CI_up_ = −0.035, *p*-value = 0.0423, Figure 4). Furthermore, lizards did not differ in their reaction towards novel food and novel objects (relative latency: MCMCglmm, estimate = −0.283, CI_low_ = −0.745, CI_up_ = 0.147, *p*-value > 0.05, Figure 4).

**Figure 4.**
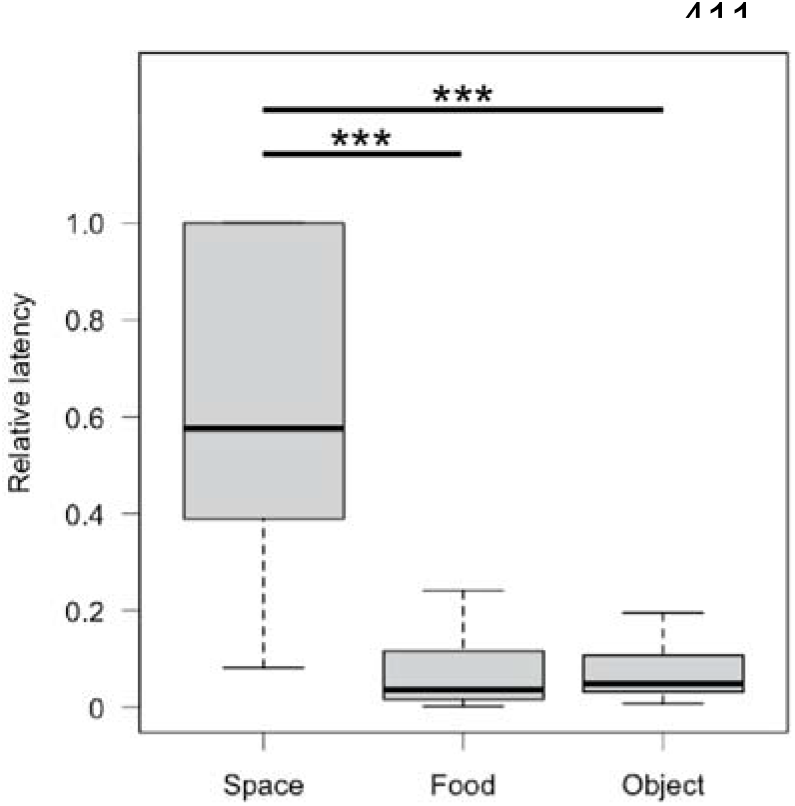
Box plot showing the relative latency to exit into a novel environment (space), to attack novel food (food) and to attack prey next to a novel object (object). The bold line indicates the median, the upper edge of the box represents the upper quartile, the lower edge the lower quartile, the top edge of the whisker the maximum and the bottom edge of the whisker the minimum (outliers are not shown). *** p < 0.001.

We found moderate evidence for a correlation between the latency to attack the novel prey item (colour pooled) and the time taken to exit a box into a novel environment (Spearman rank correlation, r_s_ = 0.573, S = 756.93, *p*-value = 0.005352; Figure 5). Latency was not correlated in any other possible combination across contexts (electronic supplementary material, Table S6 and Figure S4).

**Figure 5.**
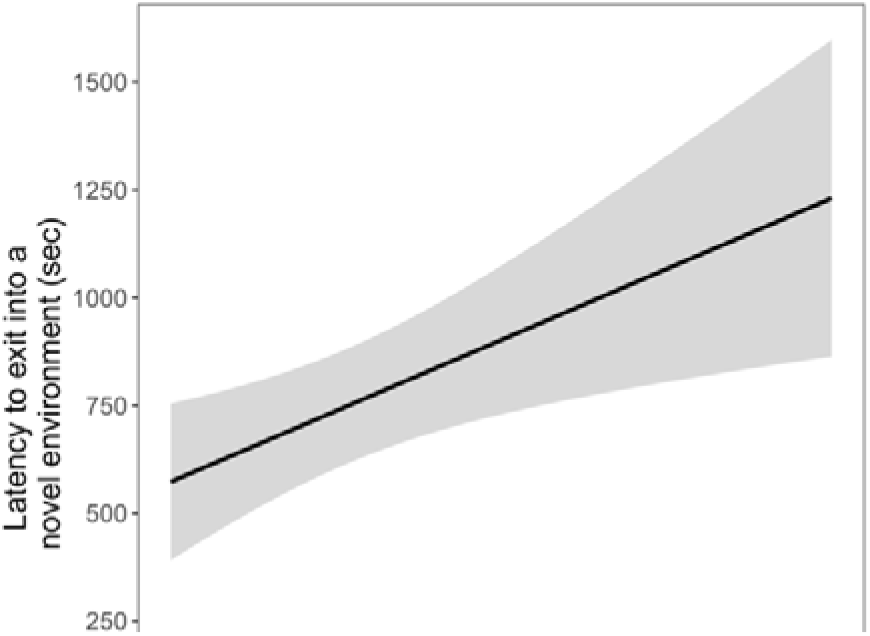
Correlation between the time to attack novel prey (regardless of colour) and to exit a box into a novel environment (capture latency). The grey area indicates the 95% confidence intervals.

We found very strong evidence that responses were repeatable across tests but at a value of only R = 0.12 (CI_low_ = 0.032, CI_up_ = 0.228, *p*-value = 0.0000255). Figure 6 shows the average latency to respond to novel stimuli across tests (object neophobia is split into responses to familiar and novel objects) for each individual. Only a few lizards were consistent across contexts. For example, lizard ID 11 showed consistently longer capture latencies, while lizard ID 14 showed consistently medium capture latencies. Contrary, lizards ID 10, 5 and 12 showed consistently shorter capture latencies across contexts.

**Figure 6.**
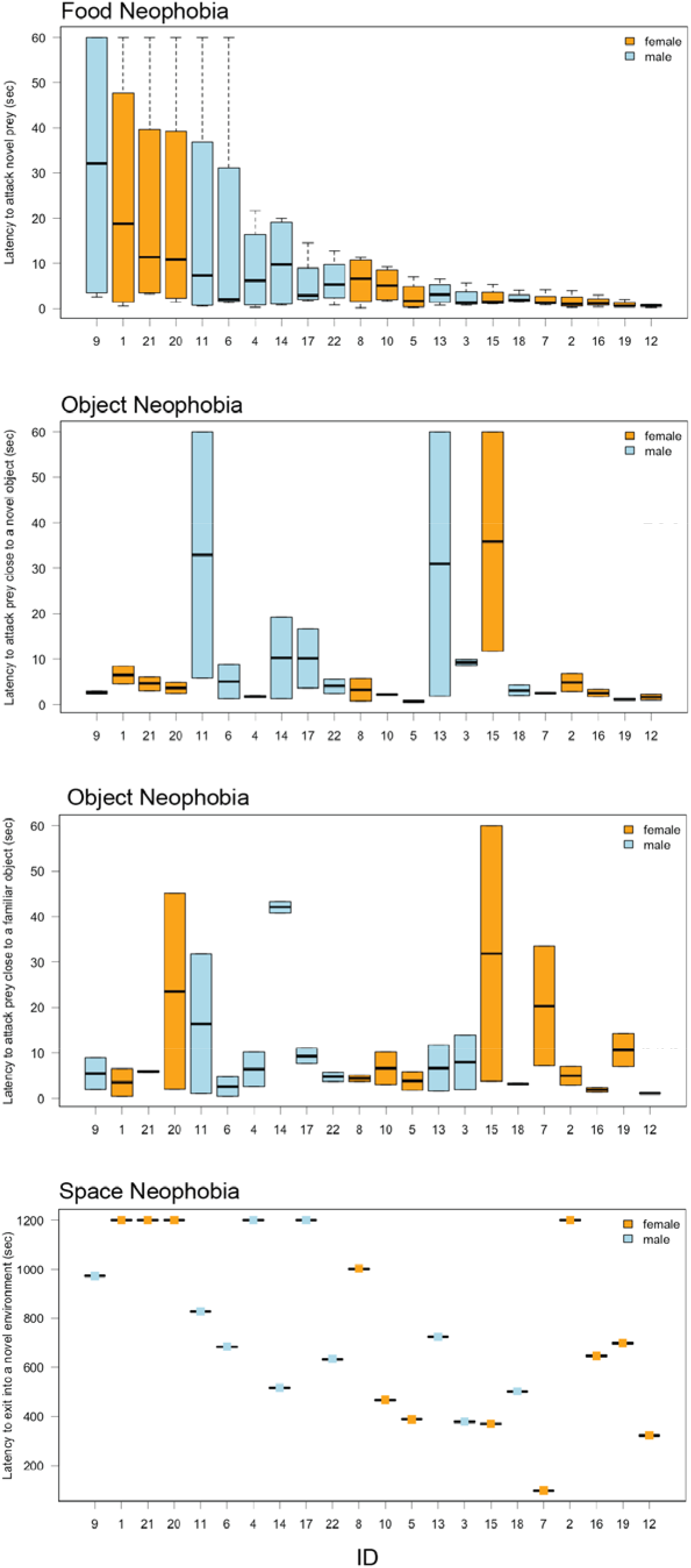
Individual capture latencies towards novel stimuli in the food, objects and space neophobia test. The x-axis shows individual IDs and is order based on the response in the food neophobia test. The bold line indicates the median, the upper edge of the box represents the upper quartile, the lower edge the lower quartile, the top edge of the whisker the maximum and the bottom edge of the whisker the minimum (outliers are not shown). Females are coloured in orange while males are coloured in blue.

## Discussion

We tested tokay geckos’ response to novelty across three contexts: novel versus familiar food (natural and coloured), familiar food and food close to familiar and novel objects as well as entering a novel space. In the food neophobia test, our geckos hesitated to attack novel food regardless of colour showing food neophobia but not dietary conservatism (Marples & Kelly, 2001). They hesitated to attack food near objects regardless of if an object was familiar or novel. We found a correlation between food neophobia and space neophobia but repeatability was low across contexts and lizards showed the strongest responses in the space neophobia test. Overall, our results demonstrate that our methodology is robust in measuring neophobia in tokay geckos.

Geckos showed the strongest responses in the space neophobia test after accounting for trial length. Such high space neophobia might evolve due to high inter- and intra-specific competition or a risky foraging environment in the wild (e.g. Elvidge, Chuard, & Brown, 2016; Brydges, Colegrave, Heathcote, & Braithwaite, 2008). Contrary, a need to exploit new habitats or migrate can lead to the evolution of low levels of space neophobia (e.g. Greenberg & Mettke-Hofmann, 2001). Little is known about the ecology of the tokay gecko in its natural habitat but adult males defend territories, females visit these territories to mate and juveniles stay with their parents after hatching until sexual maturity (Grossman, 2010). It is likely, that high levels of competition or predation pressure have led to the evolution of space neophobia in tokay geckos. Alternatively, this response might be a direct consequence of captivity in which novel space is rarely available and encountered. Future studies could explore differences in space neophobia between captive bred hatchlings, subadults ready to disperse and adults to investigate if the need to disperse leads to lower space neophobia. If no difference can be found in captivity it would point towards the lack of experience with novel space as the main cause for increased space neophobia. Furthermore, space neophobia should also be studied in wild caught individuals which would give additional evidence on if and how captivity affects space neophobia.

We found that tokay geckos hesitated to attack novel food but showed no dietary conservatism as all individuals readily consumed coloured prey and most individuals integrated the novel foods into their diet by consuming them at least on their second encounter (Marples & Kelly, 2001). Tokay geckos are dietary generalists that consume mostly insects (Aowphol, Thirakhupt, Nabhitabhata, & Voris, 2006; Grossman, 2010) but they will incorporate small vertebrates in their diet if given the opportunity (e.g. Bucol & Alcala, 2013). We would, therefore, not expect them to show dietary conservatism. In nature, tokay geckos might encounter unpalatable foods possibly leading to the evolution of some neophobic responses towards food (“dangerous niche” hypothesis; Greenberg & Mettke-Hofmann, 2001; Mettke-Hofmann et al., 2013). It is similarly likely, however, that impoverished conditions in captivity and related lack of experience with a range of prey items could explain food neophobia in our geckos and wild geckos would not show any hesitation to attack novel prey. Studies on wild individuals and investigations of colour preference or avoidance for prey in certain colours (aposematic colours such as red) will help understand if the shown hesitation is caused by evolutionary adaptation or captive conditions.

Living in high risk environments can also lead to the evolution of object neophobia (“dangerous niche” hypothesis; Greenberg & Mettke-Hofmann, 2001; Mettke-Hofmann et al., 2013). Novel objects can represent predators or dangerous foods and avoiding them could prevent death by predation or poisoning. Interestingly, familiarising our geckos with an artificial leaf for five days did not reduce neophobia. There are two reasons why geckos hesitated to feed next to a familiar and novel object. First, five days of familiarisation might not have been sufficient and in future experiments we need to familiarise lizards with the object for longer. Second, and more importantly, contrary to the cardboard objects, the leaf moved during presentation while it did not move during familiarisation. Movement of objects is rare in their captive environment as there is no strong air current for plant leaves to move. Geckos might have reacted to movement as the novel stimulus rather that the object itself. Context does affect the intensity of a neophobic response (Greenberg & Mettke-Hofmann, 2001; Ranganath & Rainer, 2003; Vernelli, 2014). A familiar but static object that suddenly moves might indicate danger. Lack of experience with natural movement might further enhance neophobic responses towards such unexpected movement which would explain the strong response we found in our object neophobia test. Researchers need to be aware of such issues to be able to interpret their animals’ responses correctly. It would be interesting to see how wild geckos with experience react in an object neophobia test comparing moving and static objects.

We found a correlation between the time taken to attack novel prey and the time taken to exit a box into a novel environment. Not many studies investigate neophobia across more than one context and there is little evidence for a correlation of neophobia across contexts (e.g. Mettke-Hofmann et al., 2002). This is also reflected in our repeatability analysis which yielded a low repeatability of only 0.12. A meta-analysis looking at repeatability in novel object tests (both neophobia and neophilia) showed an average repeatability of 0.47 (Takola et al., 2021) while another study looking at repeatability in behaviour reports an average of 0.37 (Bell, Hankison, & Laskowski, 2009). Repeatability of responses towards novel objects and novel foods in corvids was calculated around 0.5 (Miller et al., 2022). In lizards, studies have found repeatability of around 0.4 (Damas-Moreira et al., 2019; De Meester et al., 2022) but also similar low repeatability of about 0.1 (Damas-Moreira, Riley, Harris, & Whiting, 2019). Damas-Moreira and colleagues (2019) interpreted this low repeatability as evidence for flexibility. In this study, Italian wall lizards showed lower repeatability of behaviour towards novel objects compared to the Geniez’s wall lizard. Italian wall lizards are highly invasive and so are tokay geckos (Grossman, 2010; Rocha et al., 2015). Both species lower level of repeatability could possibly be an advantage when facing novel conditions during invasion into a new habitat. Alternatively, these three neophobia measures might not form a single trait but factor into different traits. Food neophobia might only be important in a foraging context while object neophobia might only be important in a predator context. More tests looking at neophobia in the context of aposematism (dangerous prey) or anti-predator neophobia could help better understand if neophobia is a single trait or not.

None of our geckos touched the novel object in the object neophilia test. Leopard geckos do investigate familiar object in new locations and unfamiliar objects in familiar locations demonstrating novelty recognition (Kundey & Phillips, 2021). This study looked at time spent close to an object rather than touching. Lizards might not readily interact with novel objects through contact. From our videos it is not possible to measure time spent close to the novel PVC cylinder because the distortion of the camera makes it impossible to determine distance from the cylinder correctly. In the future it would be beneficial to add a grid to the glass test tanks to be able to measure distance as well as exploration using our methodology. Furthermore, instead of hanging the cylinder from the center it should be place in a corner. Tokay geckos move in three-dimensional space and tests need to be adapted appropriately because all side as well as the floor and ceiling are utilised by these lizards.

Capture latency and strike latency were both good measures to detect hesitation to attack in the presence of novelty (food and object neophobia). We measured capture latency from when a lizard first noticed the food until their first attack while strike latency was measured from the last time an individual adjusted its’ gaze until the first attack. Capture latency possibly incorporates the whole decision making process including novelty recognition and decisions of how to best strike the prey for maximum capture success. Strike latency might only encompass decisions regarding capture efficiency including where to strike a novel prey item that has never been captured before or how to avoid interference of an object close to the prey. Regardless, it is more conservative to use capture latency as it is more likely to encompass the decision making process regarding novelty and should be used in future studies.

Lastly, in the food neophobia test, lizards from room 5 responded faster than lizards kept in room 2. We believe that this is caused by randomly assigning lizards to these rooms rather than any environmental factors which were identical in both rooms. By chance we might have placed a larger number of less neophobic individuals in one room than the other or more food motivated individuals in one room than the other. This is, however, of no consequence to our results because our study design ensured that each lizard acted as its’ own control.

In summary, our study reveals neophobia in different contexts in captive bred tokay geckos. Our analyses show the strongest neophobia towards entering novel space and only low repeatability and correlation across tests which might indicate that neophobia is not a single trait. Our methodology is robust providing us with the basis for future investigations into changes in neophobia related to captive conditions and rearing environment and the relationship of neophobia to different cognitive abilities.

## Supporting information

Supplementary material providing all statistical results and additional results plots

Example videos showing lizards responses to all tested stimuli

## Acknowledgements

This work was supported by the Austrian Science Fund (FWF) [grant P 31518, PI: ER] and the Swiss National Science Foundation (SNSF) [grant 197921, PI: ER].

## Notes

https://osf.io/fhw64/?view_only=bf8bef37ef7e4f55beec48706adaa82a

### Competing Interest Statement

The authors have declared no competing interest.

