## Supplementary material providing all statistical results and additional results plots for "Fear of the new? Geckos hesitate to attack novel prey, feed near objects and enter a novel space"

To

**Supplementary tables**

*Food neophobia test*

**Table S1.** Parameter estimate and test statistics of the Bayesian generalised linear mixed model analysing latency in the food neophobia test. The model included a random intercept of animal identity and a random slope of trial nested in session. The response variable latency was log transformed to fit assumptions of normality. Significant effects are highlighted in bold.

| **Parameter** | **Estimate** | **Lower 95% CI** | **Upper 95% CI** | ***p*-value** |
| --- | --- | --- | --- | --- |
| Intercept | -5.872 | -19.676 | 7.513 | 0.387846 |
| **Novel prey** | **0.625** | **0.293** | **0.951** | **0.000134** |
| Coloured prey | 0.044 | -0.286 | 0.370 | 0.790117 |
| Trial | 0.021 | -0.443 | 0.544 | 0.936361 |
| Males | -0.204 | -0.987 | 0.622 | 0.625710 |
| **Room 5** | **-0.593** | **-1.061** | **-0.095** | **0.021903** |
| Small enclosure | -0.665 | -1.447 | 0.165 | 0.096294 |
| Trial order | -0.039 | -0.194 | 0.103 | 0.608481 |
| Temperature | 0.298 | -0.254 | 0.807 | 0.263239 |
| Test order | 0.035 | -0.004 | 0.074 | 0.076127 |
| Body condition | -0.009 | -0.023 | 0.005 | 0.211018 |

**Table S2.** Parameter estimate and test statistics of the Bayesian generalised linear mixed model analysing strike time in the food neophobia test. The model included a random intercept of animal identity and a random slope of trial nested in session. The response variable latency was log transformed to fit assumptions of normality. Significant effects are highlighted in bold.

| **Parameter** | **Estimate** | **Lower 95% CI** | **Upper 95% CI** | ***p*-value** |
| --- | --- | --- | --- | --- |
| Intercept | -7.719 | -26.138 | 11.567 | 0.41095 |
| **Novel prey** | **0.503** | **0.210** | **0.796** | **0.00187** |
| Coloured prey | 0.255 | -0.038 | 0.545 | 0.08801 |
| Trial | -0.173 | -0.749 | 0.390 | 0.52741 |
| Males | -0.422 | -1.146 | 0.299 | 0.25149 |
| Room 5 | -0.341 | -0.761 | 0.074 | 0.10845 |
| Small enclosure | -0.507 | -1.209 | 0.191 | 0.14357 |
| Trial order | 0.008 | -0.131 | 0.135 | 0.90364 |
| Temperature | 0.458 | -0.075 | 1.028 | 0.10311 |
| **Test order** | **0.049** | **-0.010** | **0.088** | **0.01549** |
| Body condition | -0.004 | -0.014 | 0.007 | 0.51619 |

*Object neophobia test*

**Table S3.** Parameter estimate and test statistics of the Bayesian generalised linear mixed model analysing latency in the object neophobia test. The model included a random intercept of animal identity and a random slope of trial nested in session. The response variable latency was log transformed to fit assumptions of normality. The difference between responses towards familiar and novel objects was calculated using the posterior. Significant effects are highlighted in bold.

| **Parameter** | **Estimate** | **Lower 95% CI** | **Upper 95% CI** | ***p*-value** |
| --- | --- | --- | --- | --- |
| Intercept | 3.544 | -8.165 | 15.232 | 0.5497 |
| **Familiar object** | **1.456** | **1.050** | **1.847** | **<4e-05** |
| **Novel object** | **1.138** | **0.732** | **1.527** | **<4e-05** |
| Familiar-novel | 0.318 | -0.084 | 0.704 | > 0.05 |
| Trial | -0.189 | -0.645 | 0.260 | 0.4081 |
| Males | 0.311 | -0.472 | 1.113 | 0.4290 |
| Room 5 | -0.437 | -0.913 | 0.035 | 0.0687 |
| Small enclosure | 0.029 | -0.732 | 0.831 | 0.9346 |
| Trial order | 0.012 | -0.191 | 0.209 | 0.9048 |
| Temperature | -0.116 | -0.573 | 0.330 | 0.6155 |
| Test order | 0.012 | -0.025 | 0.048 | 0.5380 |
| Body condition | -0.003 | -0.016 | 0.010 | 0.6314 |

**Table S4.** Parameter estimate and test statistics of the Bayesian generalised linear mixed model analysing strike time in the object neophobia test. The model included a random intercept of animal identity and a random slope of trial nested in session. The response variable latency was log transformed to fit assumptions of normality. The difference between responses towards familiar and novel objects was calculated using the posterior. Significant effects are highlighted in bold.

| **Parameter** | **Estimate** | **Lower 95% CI** | **Upper 95% CI** | ***p*-value** |
| --- | --- | --- | --- | --- |
| Intercept | -0.643 | -11.352 | 9.626 | 0.901911 |
| **Familiar object** | **0.656** | **0.282** | **1.024** | **0.000437** |
| **Novel object** | **0.683** | **0.313** | **1.043** | **0.000364** |
| Familiar-novel | -0.027 | -0.395 | 0.338 | > 0.05 |
| Trial | 0.009 | -0.407 | 0.431 | 0.963421 |
| Males | **0.629** | -0.051 | 1.303 | 0.063476 |
| Room 5 | -0.308 | -0.707 | 0.085 | 0.125205 |
| **Small enclosure** | **0.971** | **0.303** | **1.639** | **0.005096** |
| Trial order | -0.054 | -0.239 | 0.128 | 0.564659 |
| Temperature | 0.009 | -0.389 | 0.414 | 0.962111 |
| Test order | 0.026 | -0.006 | 0.059 | 0.119308 |
| Body condition | -0.008 | -0.018 | 0.003 | 0.169318 |

*Space neophobia test*

**Table S5.** Parameter estimate and test statistics of the Bayesian generalised linear mixed model analysing latency in the space neophobia test. The model included a random intercept. The response variable latency was log transformed to fit assumptions of normality. Significant effects are highlighted in bold.

| **Parameter** | **Estimate** | **Lower 95% CI** | **Upper 95% CI** | ***p*-value** |
| --- | --- | --- | --- | --- |
| Intercept | -5.339 | -0.286 | 17.820 | 0.642 |
| Males | -0.129 | -1.056 | 0.854 | 0.778 |
| Room 5 | -0.243 | -0.982 | 0.448 | 0.480 |
| Small enclosure | -0.405 | -1.313 | 0.596 | 0.385 |
| Temperature | 0.496 | -0.485 | 1.468 | 0.304 |
| Test order | 0.004 | -0.049 | 0.128 | 0.334 |
| Body condition | 0.001 | -0.015 | 0.016 | 0.932 |

*Cross context correlation (syndrome)*

**Table S6.** Correlation coefficient and test statistics from the Spearman rank correlation investigating syndromes across the tested contexts (food, object and space neophobia) Significant correlations are highlighted in bold.

| **Correlation** | **rho** | **S** | ***p*-value** |
| --- | --- | --- | --- |
| Food neophobia & neophobia of a familiar object | 0.126 | 1548 | 0.5752 |
| Food neophobia & neophobia of a novel object | 0.338 | 1172 | 0.1239 |
| **Food neophobia & space neophobia** | **0.573** | **756.93** | **0.005352** |
| Neophobia of a familiar object & a novel object | 0.397 | 1068 | 0.06841 |
| Neophobia of a familiar object & space neophobia | -0.006 | 1781.1 | 0.9799 |
| Neophobia of a novel object & space neophobia | 0.193 | 1428.6 | 0.3886 |

**Supplementary figures**

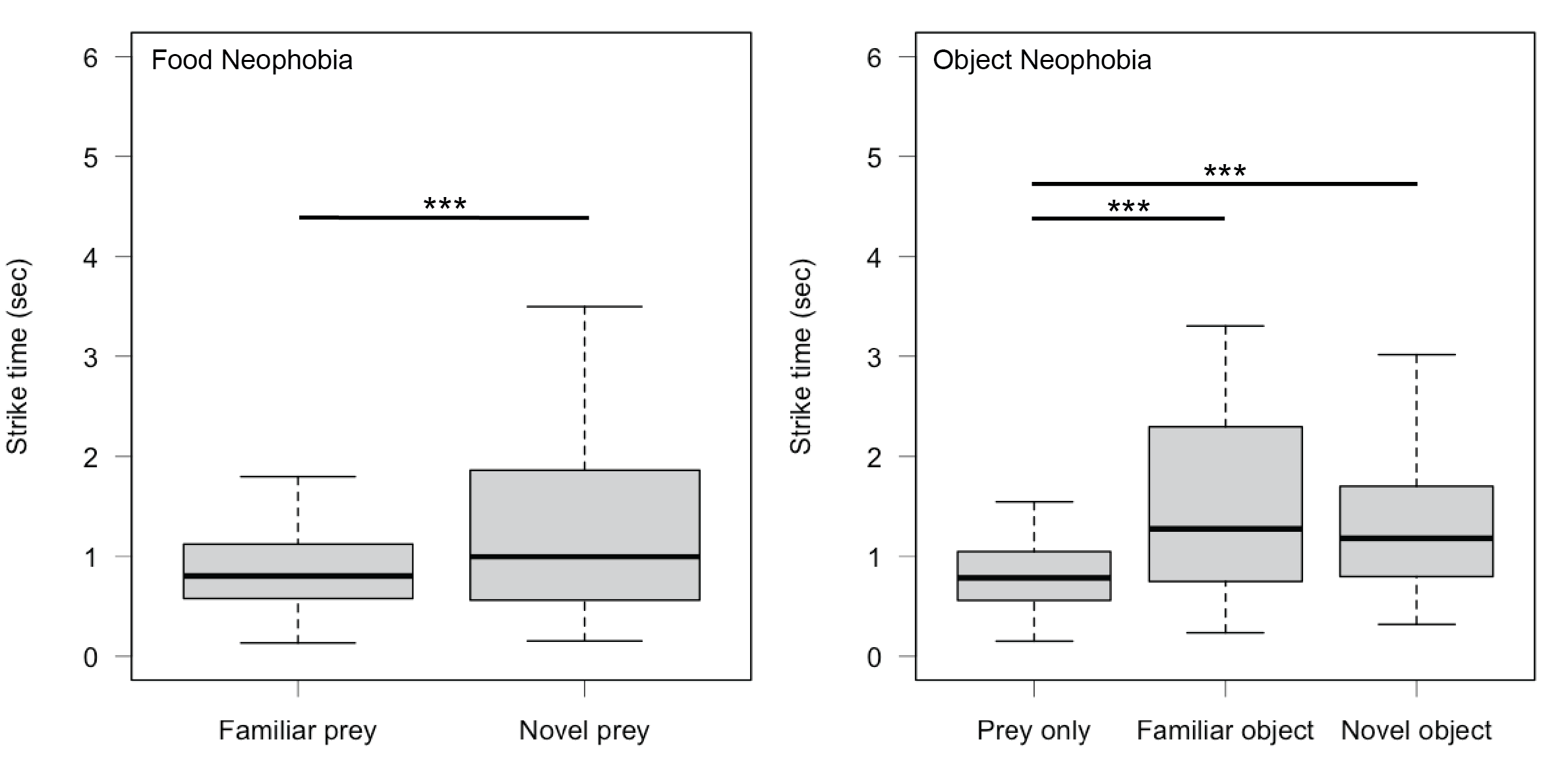

**Figure S1.** Box plots showing the strike time towards familiar (cricket) and novel food (mealworm or locust) in the food neophobia test and prey only, prey next to a familiar object (artificial leave) and a novel object (egg carton or toilet paper roll) in the object neophobia test. The bold line indicates the median, the upper edge of the box represents the upper quartile, the lower edge the lower quartile, the top edge of the whisker the maximum and the bottom edge of the whisker the minimum (outliers are not shown). *** p < 0.001.

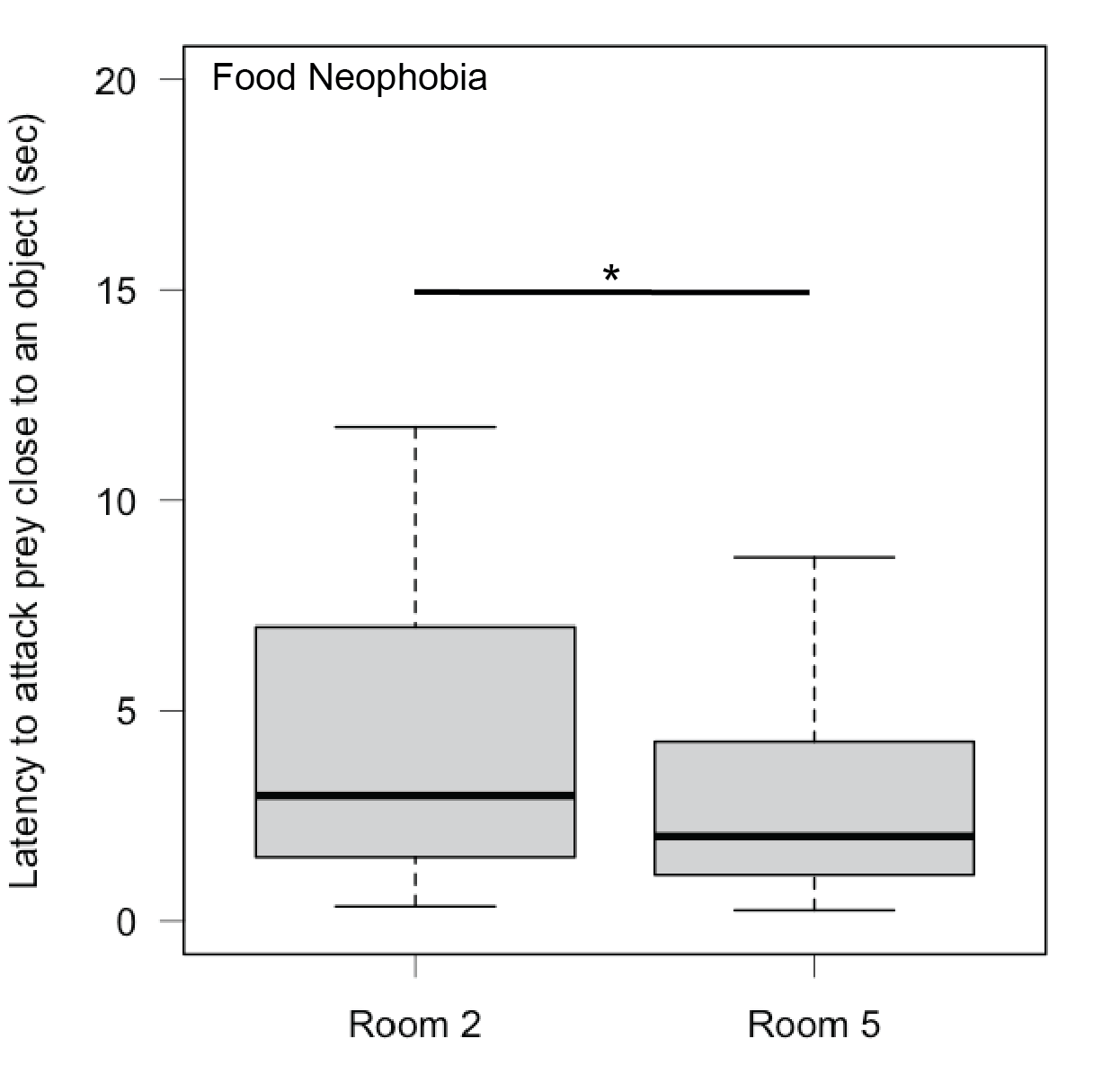

**Figure S2.** Box plots showing the latency to respond to any stimulus in individuals kept in room 2 and room 5 in the food neophobia test. The bold line indicates the median, the upper edge of the box represents the upper quartile, the lower edge the lower quartile, the top edge of the whisker the maximum and the bottom edge of the whisker the minimum (outliers are not shown). * p < 0.05.

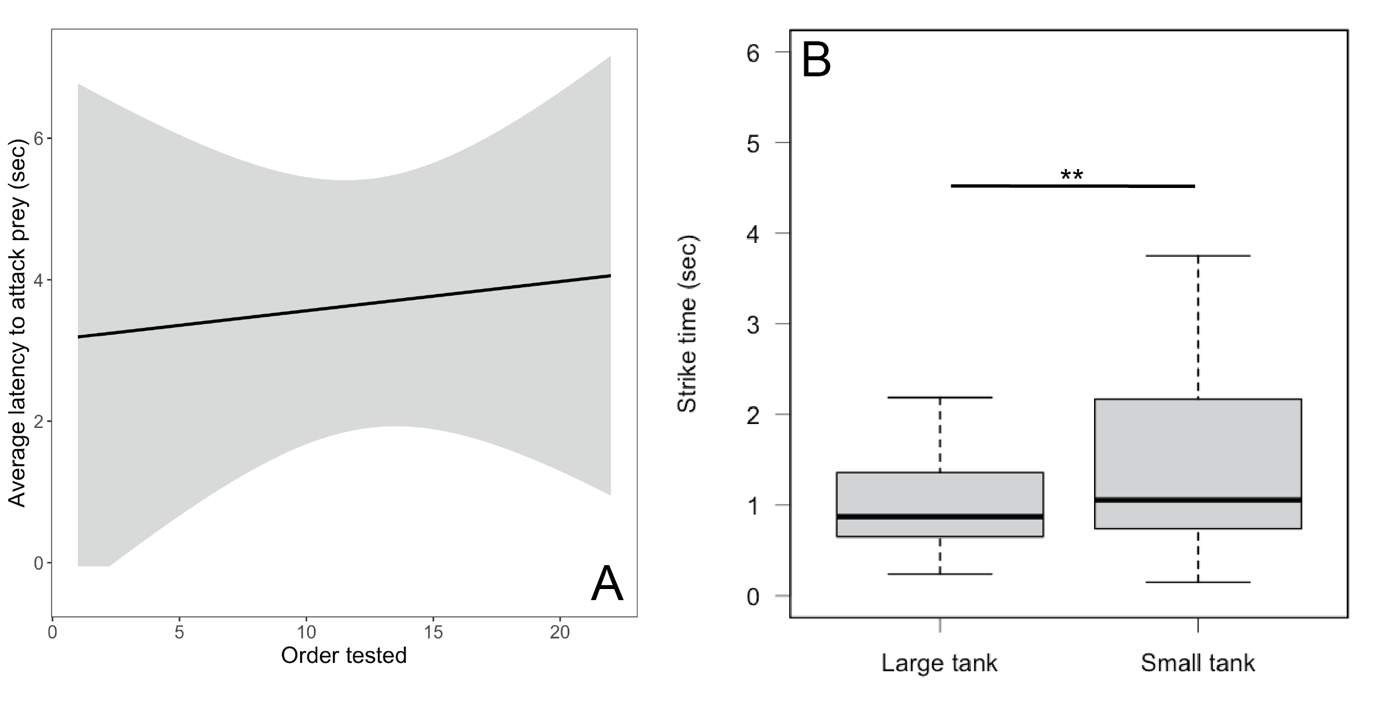

**Figure S3.** (A) Correlation between the order in which individuals were tested within a day and attack latency in the food neophobia test. The grey area indicates the 95% confidence intervals. (B) Box plot showing the difference in strike time of individuals kept in large (90x45x100 cm) and small (45x45x70 cm) tanks in the object neophobia test. The bold line indicates the median, the upper edge of the box represents the upper quartile, the lower edge the lower quartile, the top edge of the whisker the maximum and the bottom edge of the whisker the minimum (outliers are not shown). ** p < 0.01.

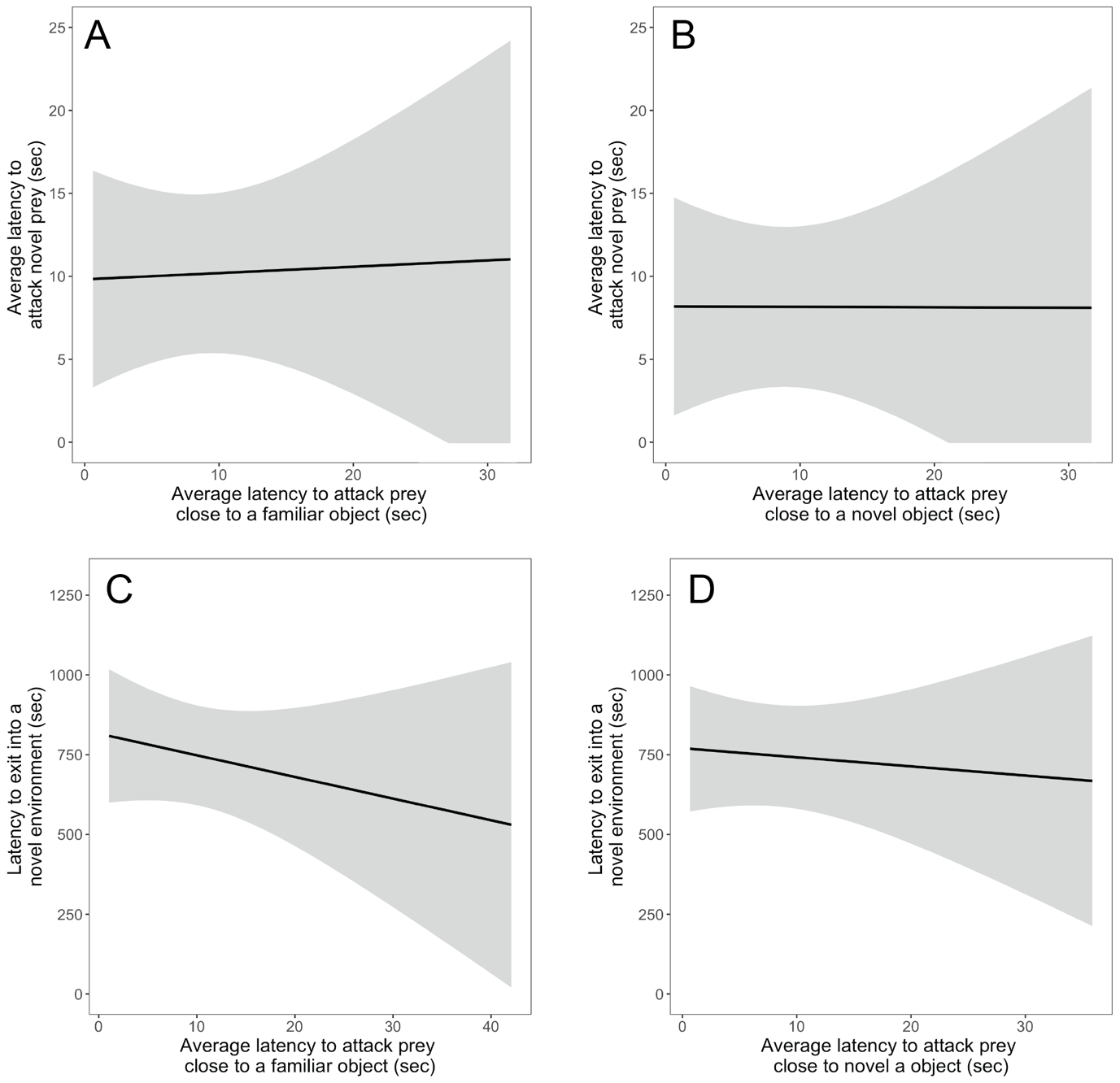

**Figure S4.** Correlation between the different contexts. (A) Correlation between the average latency to attack novel prey and the average latency to attack prey close to a familiar object. (B) Correlation between the average latency to attack novel prey and the average latency to attack prey close to a novel object. (C) Correlation between the latency to exit into a novel environment and the average latency to attack prey close to a familiar object. (D) Correlation between the latency to exit into a novel environment and the average latency to attack prey close to a novel object. The grey area indicates the 95% confidence intervals.
